# Shikimate pathway-Dependent Catabolism: enabling near-to-maximum production yield of aromatics

**DOI:** 10.1101/2024.07.06.602327

**Authors:** Lyon Bruinsma, Christos Batianis, Sara Moreno Paz, Kesi Kurnia, Job. J Dirkmaat, Alexandra Müller, Jose Juncosa Nunez, Ruud A. Weusthuis, Vitor A. P. Martins dos Santos

**Author notes:** Contributed equally (Lyon Bruinsma, Christos Batianis). Correspondence to: Christos Batianis & Vitor A. P. Martins dos Santos.

## Abstract

Catabolism is a complex network of tightly regulated metabolic reactions that provides energy and carbon to fuel anabolism in all living organisms. Rewiring catabolism is essential for harnessing industrial biotechnology but remains a substantial metabolic engineering challenge due to its high genetic stability and tight regulation acquired through evolution. In this study, by combining metabolic modeling, rational engineering, and adaptive laboratory evolution, we fundamentally redesigned bacterial catabolism. We created a new-to-nature shikimate pathway-dependent catabolism (SDC) in *Pseudomonas putida* by reprogramming the shikimate pathway as the primary catabolic route. SDC supports growth by supplying the glycerol catabolic end-product pyruvate, enabling superior production of shikimate pathway-derived molecules. Through SDC, aromatics production reached over 89% of the pathway’s maximum theoretical yield, setting a new benchmark for their microbial synthesis. Our study successfully repurposed an anabolic pathway for catabolism, exemplifying the high metabolic plasticity of microbes and providing a bacterial chassis for the efficient production of high-added value compounds.

## INTRODUCTION

Metabolism is the foundation of all organisms’ lifestyles. Despite exhibiting significant variations, virtually all living entities possess a core central carbon metabolism consisting of glycolysis, tricarboxylic acid (TCA) cycle, and the pentose phosphate pathway. This metabolic framework facilitates the conversion of carbon sources (catabolism) into crucial precursor molecules utilized in synthesizing every cellular component (anabolism) and the energy necessary for sustaining growth and maintenance^1^. Because catabolism and anabolism are highly essential for the cell, any deviation can lead to reduced cell viability or even cell death, making our view on catabolism rigid and inflexible^2^. This view hampers our ability to efficiently optimize catabolic architectures for biotechnology^3^. Extra complexity is added to these efforts, especially for anabolic pathways that are distant from catabolism, as many inherent pathways are subjected to tight regulation and therefore carry considerably low fluxes^4^.

One of these pathways is the shikimate pathway, the biochemical source of aromatic molecules including aromatic amino acids. While a myriad of valuable chemicals can be derived from this pathway, its biotechnological exploitation remains a mounting metabolic engineering challenge^5,6,7^. The shikimate pathway facilitates the biosynthesis of essential molecules, at the expense of energy use. Consequently, only a minor fraction of the assimilated carbons is directed towards this metabolic process while most are transformed via glycolysis into pyruvate that fuels the TCA cycle for generating energy. Interestingly, alongside glycolysis, organisms can produce pyruvate via various reactions, some of which pertain to the shikimate pathway. This led us to question; what would happen if cells were programmed to generate pyruvate exclusively via the shikimate pathway instead of central catabolism? Could such an approach offer possibilities for generating a synthetic catabolism aimed at chemical production rather than cellular proliferation? Although theoretically possible, implementing this unconventional metabolic configuration poses significant challenges due to the strict regulation and inflexibility of both central catabolism and the shikimate pathway. Therefore, devising a new functional catabolic system of this magnitude requires adopting smart laboratory evolutionary approaches in addition to rational engineering practices.

Recently, growth-coupled selection strategies have gained attention as an effective approach to modify cell factories for producing or consuming novel substrates or establishing new metabolic structures^8,9,10^. When employed in combination with adaptive laboratory evolution (ALE), this approach can radically reprogram cellular metabolism and yield unprecedented industrially efficient cell factories. Through ALE, microbes regain fitness by acquiring beneficial mutations that allow them to adapt to an applied selective pressure. If metabolism is artificially strained to occur via a biosynthetic pathway of interest, previously incapable of sustaining growth, evolution could enhance the flux capacity of this metabolic route, enabling so-called growth-coupled selection^11^. This approach has paved the way for installing atypical metabolisms within conventional hosts, including complete C1 assimilation by *Escherichia coli*^12,13^.

Here, through the implementation of a pyruvate-driven metabolic engineering strategy, we fundamentally restructured the catabolic network of the industrially relevant bacterium *Pseudomonas putida*, creating a Shikimate pathway-Dependent Catabolism (SDC) (Fig. 1A). By strategically disrupting central pyruvate-releasing pathways and using this pyruvate auxotroph as the base strain for ALE, we have successfully established a new glycerol catabolic pathway where pyruvate is predominantly derived from the shikimate pathway. We selected the biosynthetic pathway of 4-hydrozybenzoate (4HB) as the salvage route (Fig. 1A), as it was predicted to be the most effective compared to the other native shikimate pathway-derived pyruvate-releasing reactions. After extensive phenotypic characterization, we confirmed SDC (Fig. 1A), in which the biosynthetic pathway of 4HB acts as the primary pyruvate source. Through reverse engineering, we discovered that mutations on the regulatory genes *miaA* and *mexT* were critical in establishing this unconventional metabolic configuration. Through further model-driven metabolic engineering rounds, we achieved superior 4HB production yields in minimal medium, reaching 89% of the maximum theoretical yield. To our knowledge, this represents the highest yield ever reported. In addition, we readily adapted the SDC strain towards the production of salicylate and 3-hydroxybenzoate (3HB) with unparalleled yields. This new-to-nature catabolic architecture exemplifies the high degree of microbial metabolic plasticity and highlights the outstanding capabilities of combining rational engineering with ALE to generate new, previously unfavored, metabolisms.

**Figure 1.**
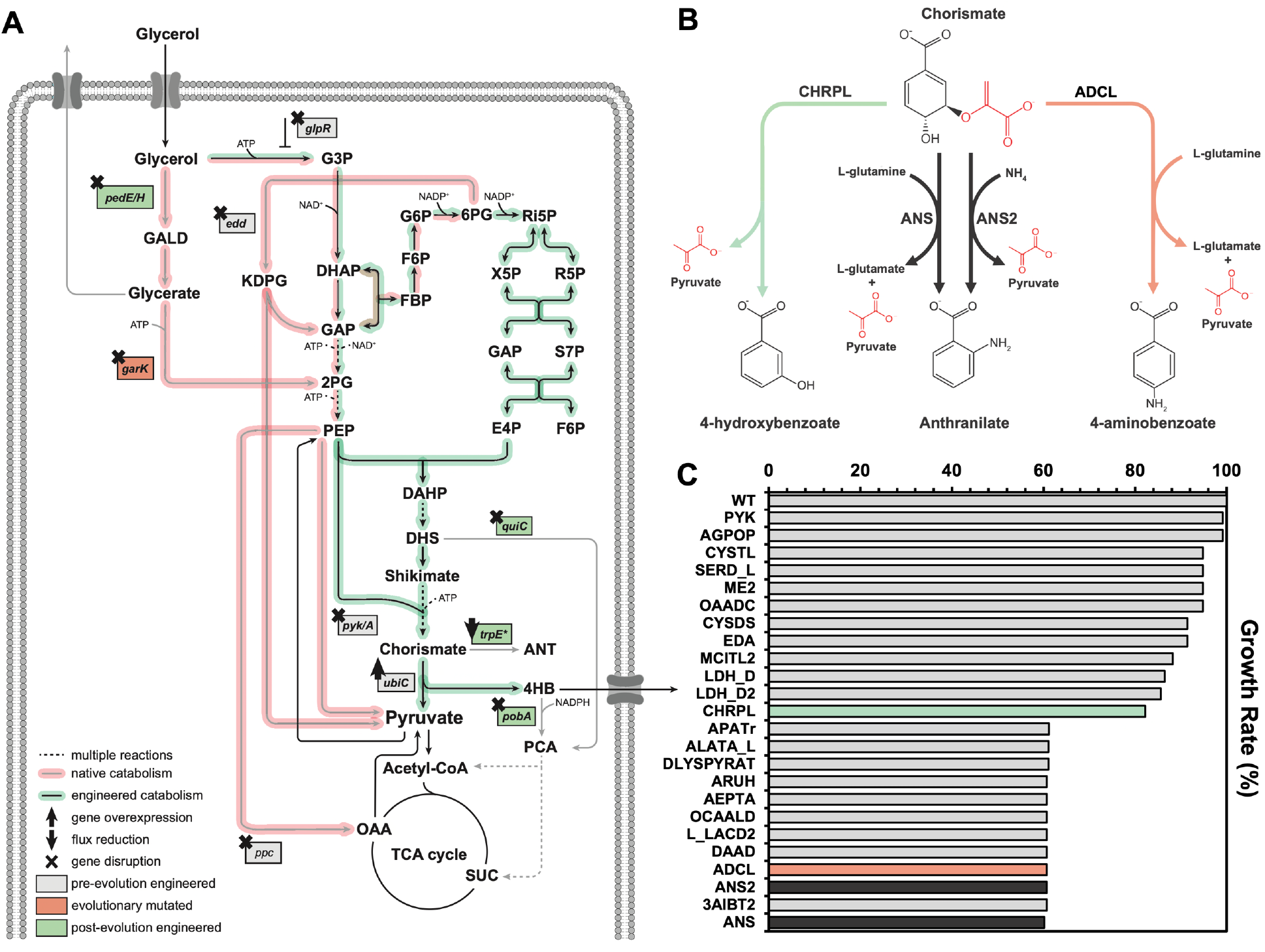
A) Metabolic architecture and mutations relevant to the Shikimate pathway-Dependent Catabolism (SDC). Abbreviations: G3P, sn-glycerol-3-P; GALD, glyceraldehyde; DHAP, dihydroxyacetone-P; GAP, glyceraldehyde-3-P; 2-PG, 2-phospho-D-glycerate; KDPG, 2-keto-3-deoxy-6-phosphogluconate; PEP, phosphoenolpyruvate; FBP, fructose-1,6-P2; F6P, fructose-6-P; G6P, glucose-6-phosphate; 6PG, 6-phosphogluconate; Ri5P, ribulose-5-P; R5P, ribose-5-P; X5P, xylose-5-P; S7P, sedoheptulose-7-P; E4P, erythrose-4-P; DAHP, 3-deoxy-d-arabinoheptulosonate-7-P; DHS, 3-dehydroshikimate; ANT, anthranilate; 4HB, 4-hydroxybenzoate; PCA, protocatechuate; SUC, succinate; OAA, oxaloacetate. B) Native shikimate-derived pyruvate-releasing reactions. C) Predicted maximum growth rates via flux balance analysis (FBA) by setting each pyruvate-releasing reaction as the sole pyruvate source. Bar colors correlate to reaction colors from panel B. Source data are provided as a Source Data file.

## RESULTS

### Metabolic design and *in-silico* assessment of SDC

Pyruvate is a key node in central metabolism predominantly produced in *P. putida* via the Entner-Doudoroff (ED) and the lower Embden–Meyerhof–Parnas (EMP) pathway^14^. Yet, apart from the main carbon metabolism, pyruvate is produced by several other innate reactions, some of which are involved in aromatic biosynthesis (Fig. 1B). We, therefore, hypothesized that one of these reactions could serve as the major pyruvate source if carbon catabolism was greatly pushed towards the shikimate pathway. To determine the most suitable reaction for promoting cell growth under SDC, we ranked all pyruvate-releasing reactions present in the iJN1463 genome-scale metabolic model using flux balance analysis (FBA). We compared the obtained *in-silico* growth rates when each of these reactions was used as the sole pyruvate source with glycerol serving as the carbon source. In all cases, growth was restored, suggesting that any of these reactions could provide adequate pyruvate for maintaining growth. Among the shikimate pathway-derived reactions (Chorismate pyruvate lyase (CHRPL), Aminobenzoate synthase (ADCL), Anthranilate synthase (ANS), Anthranilate synthase 2 (ANS2)), CHRPL was predicted to allow the highest growth rate (Fig. 1C). This reaction cleaves chorismate, the final product of the shikimate pathway, to pyruvate and 4HB (Fig. 1B). When CHRPL is the sole pyruvate source, the *in-silico* growth rate reduces by 17.8% compared to the wild-type. Still, it facilitates 26.1% faster growth compared to the other pyruvate-releasing shikimate reactions and was therefore chosen as the most efficient candidate to establish SDC.

To further assess the feasibility of SDC, the energy generation capacity of SDC and wild-type metabolism were calculated using FBA with pyruvate production as the maximization objective. While native metabolism allows the generation of 1 mol of pyruvate and 3.75 mol of ATP per mole of glycerol, SDC converts 1 mol of glycerol in 0.55 mol of pyruvate and 0.23 mol of ATP (Detailed equations in Supplementary Information). Compared to the native metabolism, SDC is energetically poor due to the need to incorporate NADPH and ATP to reach the end product chorismate. Moreover, there is a net production of CO_2_, reducing the pyruvate yield. Nonetheless, the surplus of ATP generated from reducing equivalents and available for growth and maintenance renders SDC theoretically feasible.

### Strategic design of a growth-coupled scenario for establishing SDC

Our *in-silico* analysis demonstrated the feasibility of SDC. Nevertheless, this prediction is exclusively based on stoichiometry as genome-scale metabolic models do not take genetic regulation or enzyme kinetics into consideration. At the same time, the shikimate pathway is subjected to stringent regulatory mechanisms^5^, indicating that achieving such a significant metabolic reconfiguration via rational engineering would be impracticable. Considering that, the only way to obtain such an extreme metabolic makeover is through ALE. Therefore, to install chorismate lyase as the primary pyruvate source, we created a pyruvate auxotroph to allow for pyruvate-driven evolution. This was achieved by strategically disrupting key genes in the central pyruvate biosynthesis (Fig. 2A). Initially, we deleted *edd*, encoding 6-phosphogluconate dehydratase. This reaction is part of the ED pathway together with the sequential pyruvate-releasing step, encoded by *eda*, and its deletion renders the ED pathway inactive. Next, we eliminated pyruvate kinases by deleting *pyk* and *pykA* genes to decouple the connection between pyruvate and phosphoenolpyruvate (PEP). The last major pyruvate node was removed by deleting phosphoenolpyruvate carboxylase, encoded by *ppc*. This reaction converts PEP into oxaloacetate, which subsequently can be converted to pyruvate by the oxaloacetate decarboxylase (PP_1389). It is important to note, that the deletion of *pyk, pykA*, and *ppc* not only disrupts the flux to pyruvate but also increases the intracellular PEP pool required by the shikimate pathway. Since the inactivation of the ED pathway renders *P. putida* unable to grow on glucose as a carbon source, we opted to use glycerol instead. Moreover, glycerol offers a higher maximum theoretical yield for aromatic compounds. Therefore, we deleted *glpR*, which encodes the transcriptional repressor of the glycerol catabolic operon. The final strain, termed ΔPYR, was assessed for pyruvate auxotrophy through cultivation in glycerol minimal media supplemented with increasing levels of pyruvate. As expected, ΔPYR exhibited complete pyruvate auxotrophy, as it was unable to grow without external pyruvate supplementation, with growth levels corresponding directly to the amount of added pyruvate (Fig. 2B).

**Figure 2.**
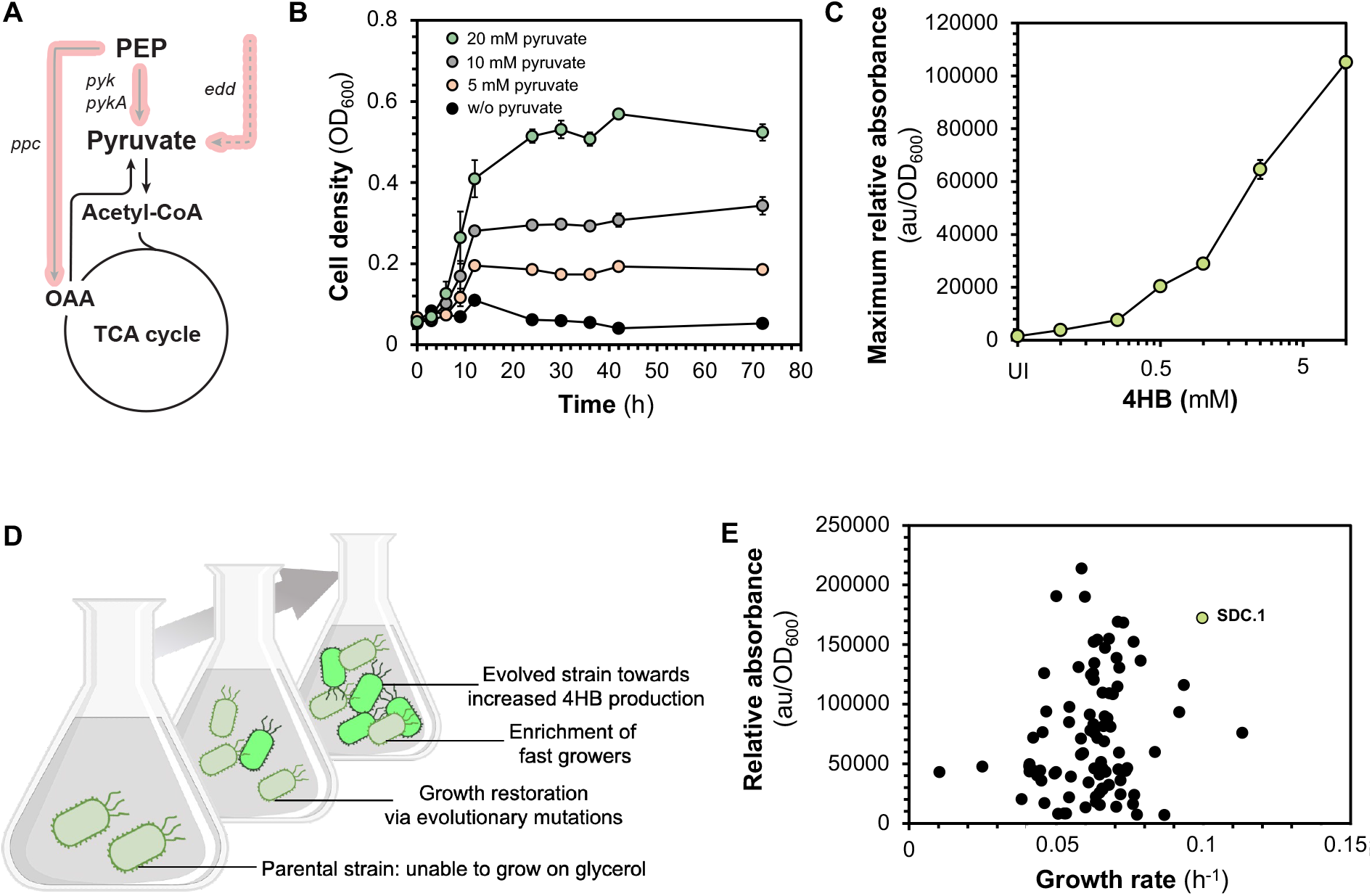
A) Metabolic scheme of the genetic basis of strain ΔPYR. All major pyruvate-producing reactions that were deleted are denoted in red. B) Growth profile of ΔPYR in glycerol minimal media supplemented with various concentrations of pyruvate. C) Dose response plot of the 4HB biosensor in *P. putida* growing under various concentrations of 4HB. D) Graphical diagram of the implemented ALE strategy. Serial passaging of strain ΔPYR carrying pSENSOR_UbiC into fresh M9 glycerol minimal medium resulting in enrichment of fast-growing phenotypes with heterogeneous fluorescence levels. E) Relative fluorescence profiles (au/OD_600_) plotted against the maximum growth rate (h^-1^) of the selected mutants after ALE in glycerol minimal medium. Every dot indicates a single isolated mutant. Error bars represent the standard deviation of three biological replicates. Source data are provided as a Source Data file.

### Implementing a second layer of screening to select superior mutants

While this auxotroph serves as a base strain for ALE, additional modifications could be necessary. Simulations revealed that alternative, more favorable, pyruvate-releasing reactions in other pathways (Fig. 1C) can allow faster growth rates than CHRPL and could be preferred during ALE. Although deleting these reactions would beneficially limit evolution towards 4HB production, this task is practically challenging as it requires the deletion of ≈20 genes, some of which could further affect cellular fitness. Notably, the other chorismate-derived pyruvate-releasing reactions (Fig. 1B) cannot be removed, as this would render the strain auxotrophic for both tryptophan and folate. Therefore, by selecting solely based on growth, there is an increased probability of identifying undesirably evolved mutants. To circumvent this problem, we implemented a second layer of screening by incorporating a previously established 4HB – responsive biosensor^15^. In this way, the desired isolates will exhibit both high growth rates and fluorescence levels after evolution (Fig. 2D). This sensor consists of the mutated transcriptional activator PobR (ΔL141, L220V) derived from *Acinetobacter bayly*i ADP1 and its corresponding promoter *P*_*pobA*_. Both genetic elements (PobR and *P*_*pobA*_), along with the *sfgfp* gene, encoding a superfolder GFP with an LAA-degradation tag, were cloned into the expression vector pSEVAb83 yielding plasmid pSENSOR (Supplementary Fig. 1A). The efficacy of the biosensor was tested in *P. putida* KT2440 Δ*glpR* growing in glycerol minimal media supplemented with increasing concentrations of 4HB. As expected, the sensor displayed dose-dependent *sfgfp* expression to exogenous 4HB concentrations (Fig. 2C, Supplementary Fig. 1B) and thus was applied as a proper selection system for the identification of evolved mutants with increased 4HB production.

Additionally, to further drive evolution towards 4HB biosynthesis, we incorporated a feedback-inhibition-resistant chorismate lyase^15^. To maintain high chorismate lyase levels, we placed the corresponding gene (*ubiC*_*fbr*_) under the control of the constitutive BBa_J23100 promoter downstream of the biosensor in the same transcriptional operon resulting in pSENSOR_UbiC (Supplementary Fig. 1A). This configuration aimed to generate a positive feedback loop, wherein 4HB production would upregulate *ubiC*_*fbr*_ expression, potentially accelerating the evolution of SDC.

### Establishing SDC via Growth-Coupled Adaptive Laboratory Evolution

We equipped *P. putida* ΔPYR with pSENSOR_UbiC and initiated evolution by cultivating the strain in glycerol minimal medium in two independent experiments. Visible growth emerged after roughly 20-24 days in both experiments (Supplementary Fig. 2). Following the breakout growth, growth became visible after only 24 h while the growth rate gradually increased as serial dilutions continued (Supplementary Fig. 2). In total, these cultures were serially diluted for ∼50 days (approximately 50 and 70 generations) and then plated on minimal media with glycerol. Forty colonies were chosen per ALE experiment based on fluorescence and selected for further characterization using a microplate reader. The selected mutants showed a diverse range of growth rates and fluorescence profiles (Fig. 2E, Supplementary Fig. 3A, B). As expected, among the mutants obtained, we observed fast-growing strains with minimal fluorescence signals that are likely to have arisen through alternative salvage pathways. This finding underscores the value of the implemented 4HB-responsive biosensor in steering the selection process after evolution. Among the diverse collection of mutants, SDC.1 emerged as a standout (Fig. 2E). This strain demonstrated fast growth among the isolated mutants (0.099 h^-1^), yet significantly slower than the control (*P. putida* Δ*glpR* carrying pSENSOR_UbiC) (0.210 h^-1^) (Supplementary Fig. 3A). However, SDC.1 displayed high levels of fluorescence compared to the other mutants and a 10-fold increase in relative fluorescence compared to the control (Fig. 2E), indicating a high flux through the shikimate pathway towards 4HB biosynthesis.

### Elucidating the genetic basis of SDC

To elucidate the genetic basis of SDC, we conducted whole-genome sequencing of the top-performing mutants. We focused solely on isolates that displayed high relative fluorescence. This is because these strains should display a high carbon flux through the shikimate pathway irrespective of the growth rate. From each evolution experiment, we chose ten mutants (full list of mutations in Supplementary Data 1). All sequenced isolates from the two individual evolution experiments contained mutations in the *miaA* and *mexT* genes at various positions (Fig. 3A). MiaA is a tRNA dimethylallyl transferase known to influence the expression of various genes related to the central and secondary metabolism^16^. In *P. putida* specifically, it was reported that the removal of *miaA*, dramatically increased the expression of the *trpE* and *trpGDC* genes, involved in tryptophan biosynthesis^17^. MiaA has been further studied in *Pseudomonas chlororaphis*, where its inactivation led to the upregulation of the *trp* genes and *aroF*^18^. The latter encodes a 3-deoxy-7-phosphoheptulonate synthase, which catalyzes the first reaction of the shikimate pathway. The *mexT* gene encodes a transcriptional regulator that has mostly been studied in *Pseudomonas aeruginosa* in which it represses the entire quinolone biosynthetic pathway and the first reaction of tryptophan biosynthesis^19^. Another major mutation that occurred in 12 out of the 20 isolates was a perturbation in *garK*, encoding a glycerate kinase. However, as the *garK* perturbation did not occur in all isolates, we deemed this one non-universal and focused solely on *miaA* and *mexT*.

**Figure 3.**
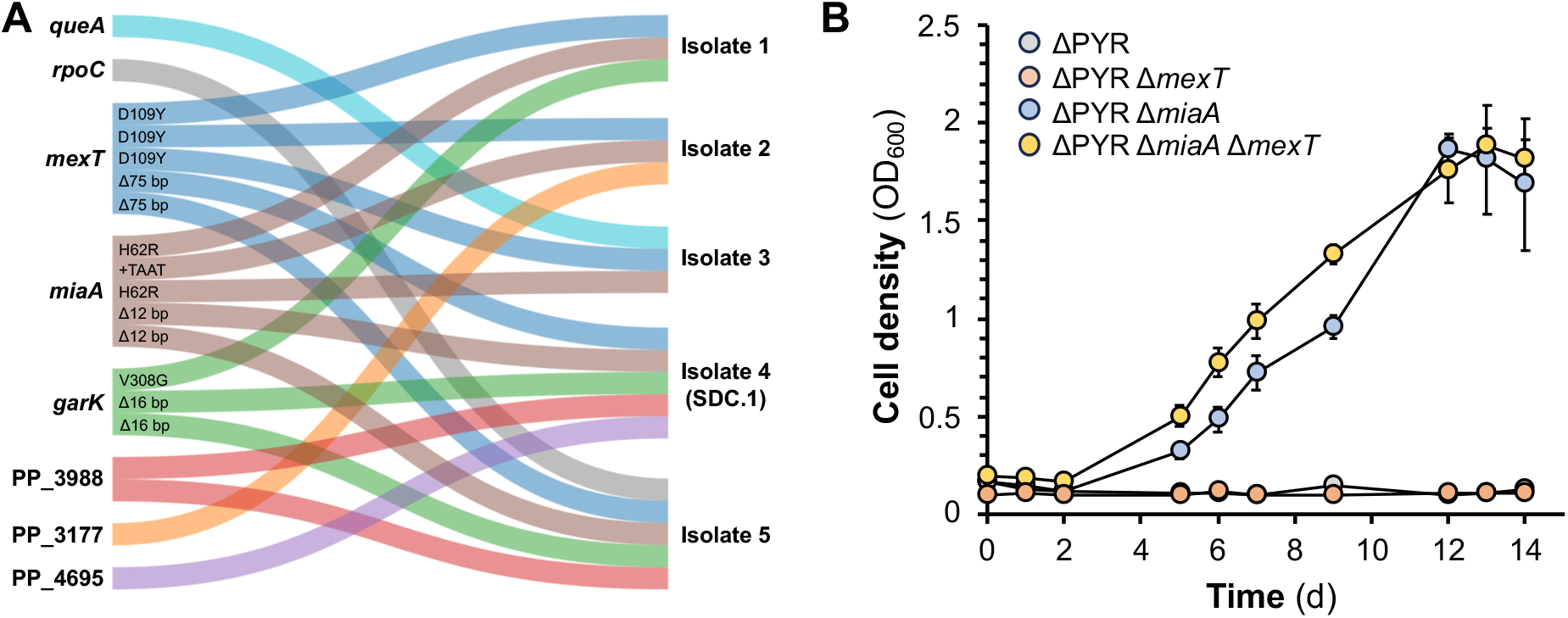
A) Major mutations identified from ALE by whole-genome sequencing of the five isolates (the Sankey diagram was built using SankeyMATIC online tool). B) Growth curves of the reverse-engineered strains. The *miaA* and *mexT* genes were mutated separately and in combination in strain ΔPYR + pSENSOR_UbiC. Genes: *queA*, S-adenosylmethionine:tRNA ribosyltransferase-isomerase; *rpoC*, DNA-directed RNA polymerase subunit beta; *mexT*, Transcriptional regulator MexT; *miaA*, tRNA dimethylallyltransferase; *garK*, glycerate kinase. Error bars represent the standard deviation of three biological replicates. Source data are provided as a Source Data file.

Moreover, in addition to genomic deletions and point mutations, sequence read coverage analysis revealed a notable amplification of transposon Tn4652 in isolate SDC.1, approximately doubling the average genomic coverage (Supplementary Fig. 4A). Tn4652 activation is recognized under stress conditions, such as starvation^20^. Upon further examination, we observed a 17kb duplication of Tn4652 within pSENSOR-UbiC, specifically in the *rep* gene, responsible for plasmid replication initiation. Subsequent analysis of the evolved plasmid highlighted a heterogeneous mixture of plasmids with varying sizes (Supplementary Fig. 4B). Evaluation following plasmid curing and transformation with the original pSENSOR-UbiC revealed comparable growth rates between strains (data not shown), suggesting that mutations in the *rep* gene may relate more to general plasmid stability during ALE^21^, rather than growth efficiency. Consequently, to mitigate plasmid heterogeneity effects and ensure the ability to retransform the plasmid upon further engineering, we opted to persist with the original plasmid.

To assess the impact of the universal mutations of *miaA and mexT*, we reverse-engineered the SDC.1 mutations of these genes into ΔPYR carrying and assessed their growth in glycerol minimal medium while expressing pSENSOR_UbiC. We selected the mutations from the SDC.1 for reverse engineering because they exhibited superior performance compared to the other isolates, as indicated by high relative fluorescence levels and growth rate. Remarkably, the *miaA* mutation alone was sufficient to permit growth in glycerol minimal medium, bypassing the need for further evolutionary adaptation (Fig. 3B). While the *mexT* mutation did not independently confer growth, its removal in combination with *miaA* led to a reduced lag phase, highlighting a synergistic effect. As the inactivation of only *miaA* allowed growth in minimal medium with glycerol in ΔPYR, we speculate that it is key in regulating the shikimate pathway and could be a potential target for metabolic engineering strategies in other organisms.

### Confirming the enhanced carbon flux through the shikimate pathway

The growth of SDC.1 was further evaluated in shake flasks containing minimal media with 40 mM of glycerol. It was confirmed that SDC.1 could grow on glycerol, albeit with a 2.2-fold slower growth rate compared to the control strain (*P. putida* Δ*glpR* carrying pSENSOR_UbiC) (Fig. 4A, B). Next, our objective was to affirm that the shikimate pathway indeed functions as the principal catabolic route for strain SDC.1. Shikimate is eventually converted to chorismate, which is subsequently cleaved into equimolar amounts of 4HB and pyruvate by chorismate pyruvate lyase. The resulting 4HB can be metabolized to protocatechuate (PCA) by p-hydroxybenzoate 3-monooxygenase, encoded by *pobA*. The produced PCA is further degraded, fueling the TCA cycle and aiding growth (Fig. 1A). Hence, we postulated that if catabolism proceeds through the shikimate pathway, deleting *pobA* will considerably impede growth and permit 4HB excretion. Additionally, using 4HB excretion as a readout, we could accurately quantify the amount of glycerol being degraded via the shikimate pathway; the higher the yield of 4HB, the more flux is diverted into this route. To test this hypothesis, we deleted the *pobA* gene in both *P. putida* Δ*glpR* and SDC.1 yielding mutants WTΔ*pobA* and SDC.2, respectively. After transformation with pSENSOR_UbiC both strains were evaluated for their ability to grow on glycerol minimal media. While the growth of WTΔ*pobA* was marginally impacted, SDC.2’s growth rate was considerably reduced by 5-fold, and it required nearly three times longer to reach the stationary phase compared to its predecessor (Fig. 4A, B). Additionally, the final cell density of SDC.2 was notably affected, showing a 1.7-fold reduction (Fig. 4A, B). This marked growth retardation in SDC.2 compared to SDC.1 provides compelling evidence of the shikimate pathway’s critical function in sustaining SDC.

**Figure 4.**
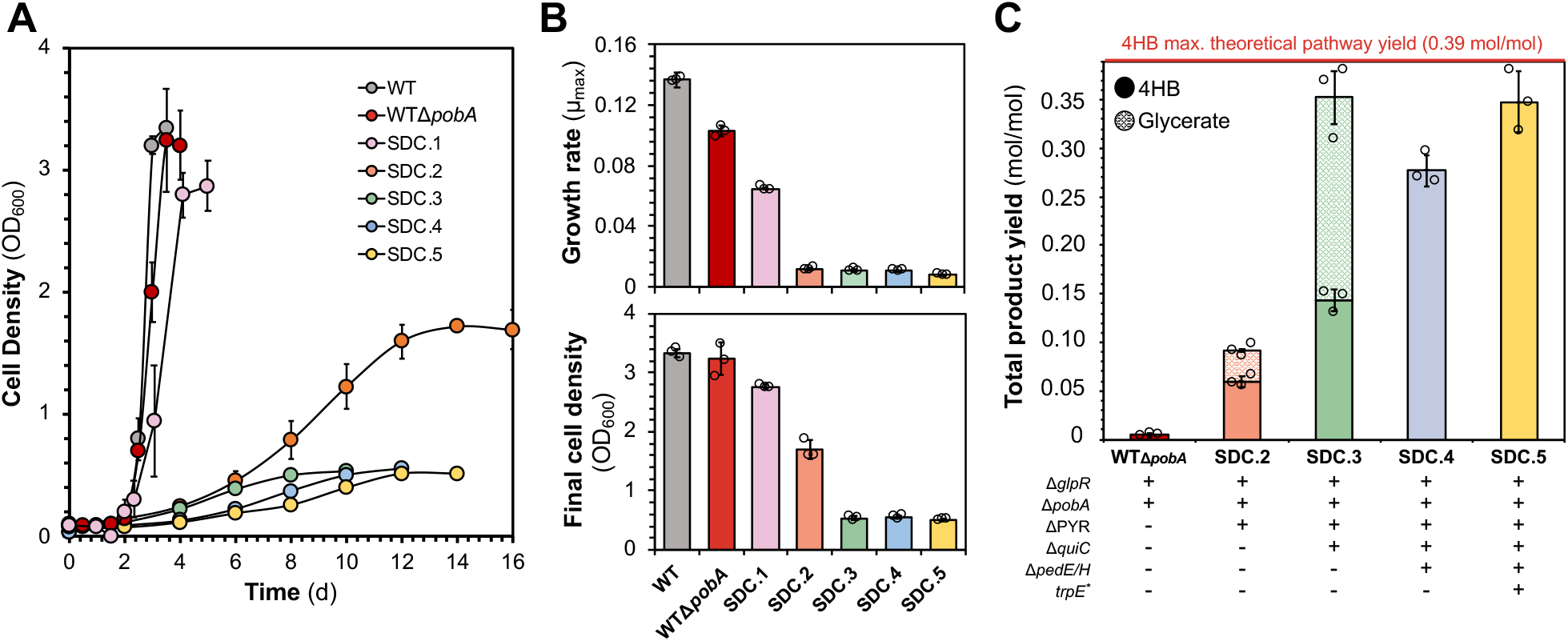
Characterization of evolved and engineered SDC strains. A) Growth curves of the WT (*P. putida* KT2440 Δ*glpR*), WTΔ*pobA* and SDC strains growing in minimal medium with 40 mM glycerol. B)Top: Maximum specific growth rates extracted from panel A. Bottom: Final cell density extracted from panel A. C) 4HB and glycerate production yields from WTΔ*pobA* and SDC variants growing in shake flasks in minimal medium with 40 mM glycerol. *trpE*_*\**_ indicates a mutated *trpE* gene (P290S). Error bars represent the standard deviation of three biological replicates. Final yields (mol/mol) and cell densities (OD_600_) have been corrected based on the quantified evaporated volume. Source data are provided as a Source Data file.

Using FBA, we predicted a maximum 4HB pathway yield of 0.39 mol/mol of glycerol. In this scenario, there is no bacterial growth, and all carbon, including the released pyruvate by CHRPL, is directed toward 4HB biosynthesis via the shikimate pathway. Therefore, we set this predicted yield as our attainable maximum representing the maximum possible flux through the shikimate pathway. The obtained 4HB yield for SDC.2 was 0.06 mol/mol, a 13.8–fold increase compared to the control strain, confirming a substantial enhancement in carbon flux through the shikimate pathway (Fig. 4C). Nonetheless, this yield accounts only for 15.4% of the predicted maximum theoretical, suggesting that considerable carbon flux remains distributed along alternative pathways. Upon simulating SDC.2 metabolism, it was predicted that 42% of the flux entering the shikimate pathway is diverted through the 3-dehydroshikimate (DHS) dehydratase (encoded by *quiC*) towards the TCA cycle (Fig. 1A, Supplementary Fig. 5). This enzyme converts the intermediate product DHS to PCA, efficiently circumventing the *pobA* deletion. To confirm this finding, we constructed SDC.3 by deleting *quiC* from SDC.2 and assessed its growth and production of 4HB. While the growth rate was slightly reduced, the most significant impact was observed in the final biomass, which decreased dramatically by 3.25-fold (Fig. 4A, B). This decrease in final biomass concentration correlated with a 2.4-fold increase in 4HB yield to attain 0.14 mol/mol (Fig. 4C). This inverse relationship between biomass formation and 4HB production emphasized the redirected metabolic flux through the shikimate pathway, demonstrating further the elevated shikimate pathway flux.

### Blocking glycerate by-production

Despite SDC.3 displaying notably high fluxes via the shikimate pathway, the 4HB yield only reached 36.7% of what was predicted as an attainable maximum. Interestingly, the initial simulations of SDC.1 predicted 19.7% of the total consumed glycerol to be excreted as glycerate (Supplementary Fig. 5). This prediction aligns with mutations in the *garK* gene among 12 out of the 20 sequenced isolates (Fig. 3A, Supplementary Data 1). SDC.1 and its derivatives contain a 16-bp deletion within this gene resulting in a frameshift. The *garK* gene encodes glycerate kinase, which is responsible for the phosphorylation of glycerate to glycerate-2-phosphate, which can then enter the main metabolism. It was therefore hypothesized that this deletion renders the enzyme inactive and allows glycerate accumulation. We quantified glycerate production in SDC.3 and its predecessor confirming that 18.3% of the total glycerol was indeed excreted as glycerate, thereby decreasing the total carbon flux towards the shikimate pathway (Fig. 4C).

According to simulations, pyruvate production from glycerol using SDC only yields 6.1% of ATP compared to the ED pathway (Supplementary Information). Aerobic bacteria like *P. putida* generate most electron carriers in the TCA cycle, which can then feed the electron transport chain and produce ATP through oxidative phosphorylation. However, in the SDC strains, the connection between glycolysis and TCA is disrupted and the TCA cycle can only be reached through the ATP-expensive shikimate pathway. Model predictions show that in native metabolism, glycerol is catabolized to the glycolytic intermediate DHAP (Supplementary Fig. 5). This process costs one ATP per glycerol and yields one reduced quinone. Alternatively, glycerol can be converted to glycerate producing two reducing equivalents that can generate ATP during oxidative phosphorylation. Natively, glycerate could then enter catabolism via its phosphorylation by GarK at the expense of ATP. Hence, it is likely that the *garK* mutation prevents ATP consumption while maintaining the generation of reduced equivalents. Consequently, in SDC.1 and its derivatives, additional ATP is generated converting glycerol into glycerate which reduces the reliance of SDC on the shikimate-mediated flux towards the TCA cycle.

To eliminate glycerate production, we eliminated the two pyrroloquinoline quinone-dependent alcohol dehydrogenases (PQQ-ADHs), PedE and PedH, responsible for converting glycerol to glyceraldehyde, resulting in SDC.4. (Fig. 1A). This deletion extended slightly the lag phase but did not have a significant impact on the growth rate (Fig. 4A, B). Glycerate production was completely abolished and revealed to be a major metabolic bottleneck as its removal increased the yield from 0.14 to 0.28 mol/mol (Fig. 4C), accounting for 71% of the maximum theoretical pathway yield.

### Reducing flux to tryptophan

To further streamline the carbon fluxes towards 4HB, we focused on branching pathways stemming from the shikimate pathway. Notable mutations in *miaA* and *mexT*, observed in the evolved strains, are known to influence tryptophan production via anthranilate synthase (ANS), encoded by *trpE* in *P. putida*^17^. Given these mutations, we hypothesized that the SDC.1 strain and its derivatives might exhibit upregulated ANS activity, providing alternative pyruvate and diverting chorismate away from 4HB synthesis. Similar to chorismate lyase, anthranilate synthase cleaves chorismate while releasing pyruvate in the process. The removal of this reaction is unwanted as it would render the strain auxotrophic to tryptophan. Therefore, we aimed to reduce its activity by introducing a P290S mutation in the *trpE* gene. This specific mutation has been reported to lower the activity of the TrpE enzyme and significantly increase the production of 4HB^22^. The new strain, termed SDC.5, exhibited a significantly slower growth rate (0.008 h^-1^) compared to SDC.4 (0.011 h^-1^) yet it attained comparable final cell densities (Fig. 3A, B). Nonetheless, the 4HB yield increased from 0.28 to 0.35 mol/mol (0.52 g/g) (Fig. 3C). This accounts for 89.2% of the maximum predicted pathway yield.

### Expanding SDC through alternative pyruvate-releasing pathways

In addition to the 4HB biosynthetic pathway, SDC has the potential to be sustained through various pyruvate-releasing reactions derived from chorismate. The shikimate pathway features several side branches that originate from chorismate and release pyruvate, including conversions of chorismate to anthranilate, 4-aminobenzoate, salicylate, 2,3-dihydroxybenzoate and 3-hydroxybenzoate (3HB) - all of which serve as precursors for a diverse range of valuable compounds^23^. To exemplify SDC’s adaptability and expand its application, we successfully integrated it with the salicylate and 3HB biosynthetic pathways (Fig. 5A).

**Figure 5.**
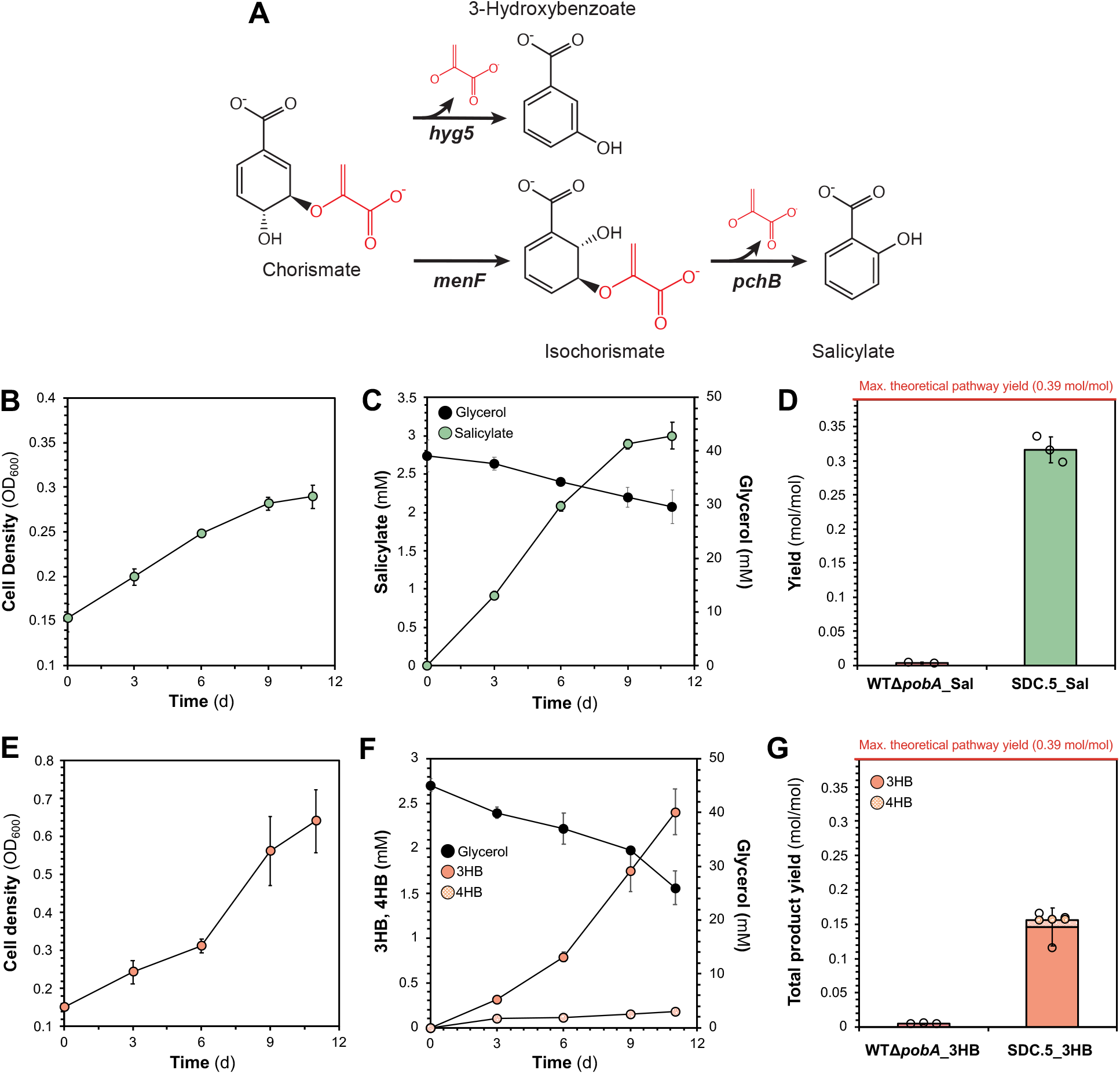
Expanded SDC towards the production of salicylate and 3HB. A) Schematic representation of salicylate and 3HB biosynthesis from chorismate. Reactions catalyzed by Hyg5 and PchB release pyruvate. B) Growth profile of SDC.5 coupled with the salicylate biosynthetic pathway. C) Salicylate production and glycerol consumption from SDC.5_sal growing in minimal medium with 40 mM glycerol. D) Salicylate production yields from WTΔ*pobA*_Sal and SDC.5_Sal growing in minimal medium with 40 mM glycerol. E) Growth profile of SDC.5 coupled with the 3HB biosynthetic pathway. F) 3HB and 4HB production and glycerol consumption from SDC.5_sal growing in minimal medium with 40 mM glycerol. G) Total product (3HB and 4HB) production yields from WTΔ*pobA*_3HB and SDC.5_3HB growing in minimal medium with 40 mM glycerol. Error bars represent the standard deviation of three biological replicates. Final yields (mol/mol) and cell densities (OD600) have been corrected based on the quantified evaporated volume. Source data are provided as a Source Data file.

Salicylate holds significant prominence as an aromatic monomer and functions as a precursor in the production of aspirin and lamivudine - an anti-HIV drug^24^. Moreover, adipic acid can be obtained from salicylate via both chemical and microbial methods, which serves as a crucial constituent for synthesizing nylon-6,625. Salicylate is synthesized from chorismate in a pathway comprising two reactions, catalyzed by the isochorismate synthase (*menF* gene product) and isochorismate pyruvate lyase (*pchB* gene product) (Fig. 5A). We placed the salicylate operon under the control of the strong inducible expression system RhaRS/P_*rhaBAD*_ in vector pSEVAb83 resulting in plasmid pSal. The strain SDC.5 was transformed with pSal yielding strain SDC.5_sal, respectively. We observed that SDC.5_sal was able to grow on glycerol demonstrating that SDC remained functional even when coupled with other pyruvate-releasing reactions (Fig. 5B). However, the growth rate and final cell density decreased by 1.7-fold compared to the SDC.5 bearing the chorismate lyase. This decrease in growth may be attributed to inadequate expression levels or catalytic speed of the corresponding enzymes. Nevertheless, the salicylate production yield reached 0.31 mol/mol, representing 81% of the maximum theoretical pathway yield (Fig. 5C, D). Although this yield represents the highest ever reported for salicylate, it falls short by about 9% when compared with what we reported previously for 4HB. However, it is important to note that in strain SDC.5_sal a flux portion is allocated towards the essential biosynthesis of 4HB thereby diminishing the total salicylate yield.

3HB is another platform chemical of industrial relevance, used as a precursor for pharmaceuticals and polymers^26,27^. Additionally, 3HB is a monomer for the production of biodegradable plastics, presenting a sustainable alternative to petroleum-based plastics^26^. Biosynthesis of 3HB occurs through the conversion of chorismate via 3-hydroxybenzoate synthase (Fig. 5A). To enable 3HB biosynthesis in SDC.5, we placed the *hyg5* gene (encoding for 3-hydroxybenzoate synthase) from *Streptomyces hygroscopicus* under the regulation of the inducible expression system RhaRS/P_*rhaBAD*_ into vector pSEVAb83, creating plasmid p3HB. The SDC.5 strain was transformed with p3HB, yielding the SDC.5_3HB strain. SDC.5_3HB demonstrated successful growth on glycerol, confirming that the SDC remains viable for this alternative pyruvate-releasing pathway (Fig. 5E). In contrast to SDC.5_sal, growth rate and final cell density appeared significantly higher, with a 2.1-fold and 2.6-fold increase, respectively. Interestingly, final cell density was also higher by 1.25-fold compared to SDC.5 expressing chorismate lyase, with both strains exhibiting identical maximum growth rates. However, this increased biomass formation resulted in significantly lower total product yield. Apart from 3HB, minor amounts of 4HB were detected as byproduct, resulting in 0.154 mol/mol total product yield, representing about 39.4% of the theoretical maximum yield (Fig 5F, G). This lower yield and increased final cell density may be attributed to potential 3HB degradation by native promiscuous enzymes that could possibly convert 3HB to PCA. Nonetheless, these results underscore the versatility of SDC, as it was proven that alternative biosynthetic pathway can sustain SDC, resulting in notably high production yields.

## DISCUSSION

Evolution has made catabolism a system of well-coordinated reactions to ensure that all living organisms obtain the maximum benefits from the available carbon resources. Unlike nature, in biotechnology organisms are programmed to utilize resources to produce valuable compounds rather than to proliferate. This is particularly challenging for anabolic pathways, such as the shikimate pathway, that are not directly connected to central metabolism, where most assimilated carbons flow.

To overcome this challenge, a new-to-nature shikimate pathway-dependent catabolism was established in *P. putida* as a universal approach for attaining high production yields of aromatic molecules. Utilizing model simulations and rational engineering, we developed a pyruvate-driven scenario for linking bacterial growth with high flux levels of the shikimate pathway. By employing evolution combined with a biosensor-assisted selection strategy, a superior mutant SDC.1 was obtained that metabolizes glycerol primarily through the shikimate pathway. Subsequently, by using a model-driven engineering approach to streamline this strain, we yielded SDC.5 which achieved 89.2% of the maximum theoretical pathway yield for 4HB; thus, demonstrating significant fluxes throughout the shikimate pathway.

Growth-coupled bioproduction of shikimate-derived products using pyruvate-releasing reactions has been theorized before^5^ and has recently been proven^28,29^. Nevertheless, these approaches still necessitate the external provision of aromatic amino acids and yeast extract indicating a deficient shikimate-pathway flux to facilitate proliferation. The intricate regulation of the shikimate pathway may be responsible for the auxotrophic nature observed in these designs, which is caused by inadequate carbon flux. This pathway is controlled through multiple means such as transcriptional repression, attenuation, and feedback inhibition7, posing a challenge for conventional metabolic engineering techniques. Our findings demonstrate that regulation represents the primary bottleneck encountered. The establishment of SDC was facilitated by only a few mutations that likely disrupted the regulatory network governing the shikimate pathway. Genetic alterations were detected across all sequenced isolates within *miaA* and *mexT* genes; both of which encode proteins involved in regulation, although their exact mechanisms remain elusive. Elimination of *miaA* in strain ΔPYR enabled growth restoration and subsequent elimination of *mexT* led to further enhancements in growth. Thus, it is worthwhile exploring these genes as potential metabolic engineering targets for the shikimate pathway within other industrial organisms.

Metabolic modeling was frequently used to guide our engineering efforts. One notable example is the prediction of glycerate as a by-product. As SDC is energetically poor, accumulation of glycerate likely occurred to conserve ATP. Moreover, glycerate production results in the generation of reduced cofactors which can be used for respiration, decreasing the required catabolic flux over the shikimate pathway. In general, ATP seems to be the most limiting factor in the fitness of SDC. The pyruvate released by chorismate lyase can be oxidized in the TCA cycle to generate reducing equivalents for respiration. However, the high yields observed in the SDC.5 strain show the recycling of pyruvate to PEP to further fuel 4HB production. Nevertheless, this step also requires the incorporation of ATP. In SDC, this behaviour would promote product formation but results in low growth rates that could limit productivity. Therefore, to increase the ATP pool in SDC, external electron donors could be applied to generate NADH, which then can be oxidized in the electron transport chain to provide ATP. A potential electron donor is phosphite, whose potential has recently been explored in *P. putida*^30^. Although its dehydrogenase is kinetically slower than its formate counterpart^31^, it was able to increase the NADH pool in *P. putida*. Moreover, compared to formate, phosphite metabolism gives a competitive advantage, allowing non-sterile fermentation conditions, and lowering overall production costs^31,32^. Therefore, both electron donors are worthy of investigation to further improve the SDC.

To further realize the full potential of the SDC, productivity optimization is an unavoidable challenge. Although the yield was significantly increased, productivity is still limited in SDC.5. In our current design, the shikimate pathway carries the whole metabolic regime with only a single overexpression. Thus, further optimization would be a necessity to increase the fluxes and improve productivity. This could either be achieved through genomic overexpression of key enzymes or a second round of laboratory evolution. The streamlined SDC.5 strain has a maximized flux through the shikimate pathway where the released pyruvate is the only junction between glycerol catabolism and the TCA cycle. Therefore, this strain would be an excellent candidate for another round of evolution. Additionally, fluxes in SDC could be optimized through rational, model-guided engineering. We demonstrated the power of metabolic modeling to aid the experimental design, identify metabolic bottlenecks, and guide optimization based on maximum yields and optimal growth rates. Compared to the wild-type, SDC strains have a complicated glycerol-to-pyruvate node with more enzymatic steps that make rational engineering nontrivial. For an optimal flux, a fine balance in erythrose-4-phosphate (E4P) and phosphoenolpyruvate (PEP) is required. We hypothesize that the main limitation of SDC is the availability of E4P, as the PEP pool is already significantly increased through the blockage of its degradation nodes in the initial ΔPYR strain. Like the shikimate pathway, the pentose phosphate pathway is predominantly used for anabolic reactions in *P. putida*, likely displaying low native fluxes^33^. However, to fully realize the potential of metabolic modeling, more information regarding the metabolic fluxes and regulatory organization would need to be gathered. All this information could be fed back into the model to select overexpression targets that, while maximizing growth, minimize resource allocation towards the expression of unnecessary enzymes and accelerate the establishment of a superior SDC strain with high yields and productivity.

This work highlights the plasticity of bacterial metabolism and how a combination of model-driven design, rational engineering, and laboratory evolution can create novel metabolisms. We repurposed the shikimate pathway as the major catabolic route and demonstrate that it can carry the whole cellular flux for growth in a versatile way by just exchanging the final pyruvate-releasing reaction. We therefore envision the SDC strain as a platform plug-in strain for the high-yield production of a myriad of aromatics with minimal engineering efforts.

## MATERIALS AND METHODS

### Plasmids, primers, and strains

All strains and plasmids used in the present study are listed in Supplementary Data 2. Primers used for plasmid construction and gene deletions are listed in Supplementary Data 3.

### Culture conditions

*P. putida* and *E. coli* cultures were incubated at 30^°^C and 37^°^C respectively. For overnight cultures, both strains were propagated in Lysogeny Broth (LB) medium containing 10 g/L NaCl, 10 g/L tryptone, and 5 g/L yeast extract. For the preparation of solid media, 1.5% (w/v) agar was added. Antibiotics, when required, were used at the following concentrations: kanamycin (Km) 50 μg/ml, gentamycin (Gm) 15 μg/ml, chloramphenicol (Cm) 50 μg/ml and apramycin (Apra) 50 μg/ml. All growth and production experiments were performed using M9 minimal medium (per Liter; 3.88 g K_2_HPO_4_, 1.63 g NaH_2_PO_4_, 2.0 g (NH_4_)_2_SO_4_, pH 7.0). The M9 media was supplemented with a trace elements solution (10 mg/L ethylenediaminetetraacetic acid (EDTA), 0.1 g/L MgCl_2_·6H_2_O, 2 mg/L ZnSO_4_·7H_2_O, 1 mg/L CaCl_2_·2H_2_O, 5 mg/L FeSO_4_·7H_2_O, 0.2 mg/L Na_2_MoO_4_·2H_2_O, 0.2 mg/L CuSO_4_·5H_2_O, 0.4 mg/L CoCl_2_·6H_2_O, 1 mg/L MnCl_2_·2H_2_O). In these experiments, strains were precultured in 10 ml LB with corresponding antibiotics. Then, the cultures were washed twice in M9 media without a carbon source. Finally, the cultures were diluted to an OD_600_ of 0.1 to start the experiment. Flask experiments were performed in 250 ml Erlenmeyer flasks filled with 25 ml of M9 minimal medium with 40 mM (SDC characterization) or 200 mM (ALE experiment) glycerol and the cultures were incubated in a rotary shaker at 200 rpm at 30^°^C.

### Cloning procedures

Plasmids were constructed using the standard protocols of the previously described SevaBrick assembly^34^. All DNA fragments were amplified using Q5^®^ Hot Start High-Fidelity DNA Polymerase (New England Biolabs). To construct the plasmids pSENSOR and pSENSOR_UbiC, the 4HB-responsive regulator and promoter (PobR/*P*_*pobA*_) were synthesized by Integrated DNA Technologies (IDT), and the *ubiC*^*E31Q*/*M34V*^ was amplified from the genome of *E. coli* with primers to introduce the E31Q and M34V mutations. All parts were integrated into the pSB1C3 repository and subsequently assembled into pSEVAb83 following the standard procedures of the SevaBrick assembly. For the construction of the pSal plasmid, the *menF* and *pchB* genes were amplified from *E. coli* and *Pseudomonas aeruginosa*, respectively, using colony PCR. Genes were domesticated in the repository vector pSB1C3 and subsequently cloned into a pre-linearized pSEVAb83 backbone under the control of the inducible expression system RhaRS/P_*rhaBAD*_ by SevaBrick assembly. For the construction of the p3HB plasmid, the *hyg5* gene from *Streptomyces hygroscopicus* was synthesized by Integrated DNA Technologies (IDT) and cloned under the control of the inducible expression system RhaRS/P_*rhaBAD*_ into pSEVAb83 backbone using the same procedures. All plasmids were transformed via heat shock in chemically competent *E. coli* DH5α λpyr cells and via electroporation or conjugation in *P. putida*. Transformants were selected on LB agar plates with corresponding antibiotics and colonies were tested by colony PCR with Phire Hot Start II DNA polymerase (Thermo Fisher Scientific). After extraction, all constructs were verified by Sanger DNA sequencing (MACROGEN Inc.).

### Genome modifications

Genomic modifications in this study were performed using the protocols previously described by Wirth et al^35^ and Volke et al^36^. Homology regions of ± 500 bp were amplified up and downstream of the target gene from the genome of *P. putida* KT2440. Both regions were cloned into the non-replicative pGNW vector and propagated in *E. coli* DH5α λpir. Correct plasmids were transformed into *P. putida* by electroporation and selected on LB + Km plates. Successful co-integrations were verified by PCR. Hereafter, co-integrated strains were transformed with the pQURE6-H, and transformants were plated on LB + Gm containing 2 mM 3-methylbenzoic acid (3-mBz). This compound induces the XylS – dependent *Pm* promoter, regulating the *I-Sce*I homing nuclease that cuts the integrated pGNW vector. Successful gene deletions were verified by PCR and Sanger sequencing (MACROGEN inc). The P290S modification in *trpE* was verified by MASC PCR as described by Asin-Garcia et al^37^. Hereafter, the pQURE6-H was cured by removing the selective pressure and its loss was verified by sensitivity to gentamycin.

### Analytical methods

Cell growth was determined by measuring the optical density at 600 nm (OD_600_) using an OD600 DiluPhotometer spectrophotometer (IMPLEN) or a Synergy plate reader (BioTek Instruments). Analysis of glycerol and pyruvate in supernatants was performed using high-performance liquid chromatography (HPLC) (Thermo Fisher Scientific) equipped with an Aminex HPX-87H column. The mobile phase was 5 mM of H_2_SO_4_ at a flow rate of 0.6 ml/min, the column temperatures were held at 60 °C and the compounds were detected using a Shodex RI-101 detector (Shodex). The amount of produced 4HB was determined using HPLC (Shimadzu) with a C18 column (4.6 mm × 250 mm) and a UV/vis detector set at 472 nm. The mobile phase consisted of Milli-Q water (A), 100 mM formic acid (B), and acetonitrile (C) with a flow rate of 1 ml/min at 30 °C. Chromatographic separation of analytes was attained using the following gradient program: *t* = 0 – 5 min: A-55%, B-10%, and C-35%; from *t* = 5 – 10 min ramp to A-10%, B-10%, and C-80% and held until 15 min. Then from *t* = 15 – 16 min, the gradient was returned to A-55%, B-10%, and C-35% and maintained isocratic for a total run time of 18 min. For quantification, calibration curves were prepared using pure standards (99% purity) purchased from Sigma-Aldrich.

### Adaptive laboratory evolution

Strain ΔPYR was inoculated in two 250 ml shake flasks with M9 minimal medium supplemented with 200 mM glycerol as the sole carbon source and incubated in a rotary shaker at 200 rpm at 30 ^°^C. The starting cell density was set at OD_600_ = 0.1 and cells were diluted back to the same OD_600_ after they reached an OD_600_ of >1.0. Evolved strains were selected on M9 agar plates with 200 mM glycerol and characterized in 200 μl of M9 medium with 40 mM glycerol using a Synergy plate reader (BioTek Instruments). Cell density (OD_600_) and GFP fluorescence (excitation 485 nm, emission 512 nm, gain 50) were measured over time using continuous linear shaking (567 cpm, 3mm), and measurements were taken every fifteen minutes. The number of generations was calculated based on the OD_600_ value, according to equation:

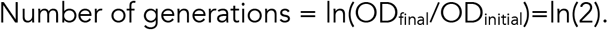

A customized Python script was programmed to calculate the growth rates taking the natural log of the OD_600_ values.

### Whole-genome sequencing

The genomic DNA of the ΔPYR and evolved mutants was isolated from LB overnight cultures using the GenElute™ Bacterial Genomic DNA Kit (Sigma-Aldrich St. Louis, MO). The extracted DNA was evaluated by gel electrophoresis and quantified by a NanoDrop spectrophotometer (Thermo Fisher Scientific). Samples were sent for Illumina sequencing to Novogene Co. Ltd. (Beijing, China). Raw Illumina reads were trimmed for low quality and adapters with fastp (v0.20.0). Mutations were identified by comparing the reads to the annotated reference genome of *Pseudomonas putida* KT2440 (GCF_000007565.2) using breseq (v0.35.5)^38^.

### Genome-scale metabolic modeling

Computational analysis was performed using COBRApy (version 0.18.1) and Python (version 3.6, Python Software Foundation). We used iJN1462, the latest developed genome-scale model (GEM) of *P. putida* to rank pyruvate-releasing reactions and to simulate native and SDC metabolism^39^. In all simulations, glycerol was used as the sole carbon source with a maximum uptake rate of 3.95 mmol/g_CDW_/h^40^. All metabolic reactions able to produce pyruvate in the iJN1462 GEM were evaluated for their capacity to support growth as the sole pyruvate source. The upper and lower bounds of all pyruvate-releasing reactions were constrained to zero except for two essential reactions: ANS2 (*trpE, pabA*) and ADCL (*pabC*). Iteratively, the flux through each reaction was unconstrained making it the main available source of pyruvate in the model. The growth rate, represented by the BIOMASS_KT2440_WT3 reaction, was maximized. Reactions were ranked according to the predicted maximum growth rates relative to the wild-type growth rate (100%). iJN1462 was modified to correctly simulate SDC and wild-type metabolism^41^ (Supplementary Data 4). When simulating SDC, a pFBA-like constraint was applied such that the sum of all the predicted fluxes cannot exceed the sum of all predicted fluxes in native metabolism. Besides, as a base for SDC simulation, we constrained to zero the flux through reactions EDD, PYK, AGPOP, ME2, OAAFC, SERD_L, CYSTL, CYSDS, MCITL2, LDH_2, and LDH_D2 to reproduce the result of the biosensor-assisted ALE that made CHRPL the main pyruvate source. Model modifications that allowed the simulation of the different SDC strains are presented in Supplementary Data 5. Maximum theoretical 4HB yields were calculated maximizing the 4HB exchange reaction (EX_4hbz_e). The optimal metabolism of the SDC strains and their maximum growth rates were simulated maximizing biomass production (BIOMASS_KT2440_WT3).

## ACKNOWLEDGEMENTS

This project was funded by the European Union Horizon2020 projects EmPowerPutida (grant number 635536), IBISBA (grant numbers 730976 and 871118) and BIOS (grant number 101070281) and the NWO (project number GSGT.2019.008).

## AUTHOR CONTRIBUTIONS

C.B. conceived this study. C.B. and L.B. designed the study. L.B., C.B., K.K., J.J.D., A.M., and J.J.N. conducted the experiments. S.P.M. conducted the computations. V.A.P.M.d.S. and R.A.W. supervised this study. V.A.P.M.d.S. arranged funding.

## REFERENCES

1. Noor, E., Eden, E., Milo, R. & Alon, U. Central carbon metabolism as a minimal biochemical walk between precursors for biomass and energy. Mol. Cell. 39.5, 809–820 (2010).

2. Stephanopoulos, G, & J J Vallino. Network rigidity and metabolic engineering in metabolite overproduction. Science 252, 5013 (1991).

3. Staffas, L., Gustavsson, M., & McCormick, K. Strategies and policies for the bioeconomy and bio-based economy: An analysis of official national approaches. Sustainability 5, 2751–2769 (2013).

4. Nielsen, J., & Keasling, J. D. Engineering Cellular Metabolism. Cell 164, 1185–1197 (2016).

5. Averesch, N. J. H., & Krömer, J. O. Metabolic engineering of the shikimate pathway for production of aromatics and derived compounds—present and future strain construction strategies. Front. Bioeng. Biotechnol. 6, 32 (2018).

6. Fujiwara, R., Noda, S., Tanaka, T., & Kondo, A. Metabolic engineering of Escherichia coli for shikimate pathway derivative production from glucose–xylose co-substrate. Nat. Commun. 11, 279 (2020).

7. Li, M., Liu, C., Yang, J., Nian, R., Xian, M., Li, F., & Zhang, H. Common problems associated with the microbial productions of aromatic compounds and corresponding metabolic engineering strategies. Biotechnol. Adv. 41, 107548 (2020).

8. Kim, S. et al. Growth of E. coli on formate and methanol via the reductive glycine pathway. Nat. Chem. Biol. 16, 538–545 (2020).

9. Nielsen, J. R., Weusthuis, R. A., & Huang, W. E. Growth-coupled enzyme engineering through manipulation of redox cofactor regeneration. Biotechnol. Adv. 63.9, (2023).

10. Orsi, E., Claassens, N. J., Nikel, P. I., & Lindner, S. N. Growth-coupled selection of synthetic modules to accelerate cell factory development. Nat. Commun. 12.1, 1–5. (2021).

11. Iacometti, C., Marx, K., Hönick, M., Biletskaia, V., Schulz-Mirbach, H., Dronsella, B., & Lindner, S. N. Activating silent glycolysis bypasses in Escherichia coli. Biodes. Res. (2022).

12. Gleizer, S. et al. Conversion of Escherichia coli to generate all biomass carbon from CO2. Cell 179.6, 1255–1263 (2019).

13. Chen, F. Y. H., Jung, H. W., Tsuei, C. Y., & Liao, J. C. Converting Escherichia coli to a synthetic methylotroph growing solely on methanol. Cell 182, 933–946 (2020).

14. Nikel, P. I., Chavarria, M., Fuhrer, T., Sauer, U. & de Lorenzo, V. Pseudomonas putida KT2440 strain metabolizes glucose through a cycle formed by enzymes of the Entner Doudoroff, Embden-Meyerhof-Parnas, and pentose phosphate pathways. J. Biol. Chem. 290, 25920–25932 (2015).

15. Jha, R. K. et al. Sensor-enabled alleviation of product inhibition in chorismate pyruvate-lyase. Acs. Synth. Biol. 8, 775–786 (2019).

16. Koshla, O. et al. Gene miaA for post-transcriptional modification of tRNAXXA is important for morphological and metabolic differentiation in Streptomyces. Mol. Microb. 112, 249–265 (2019).

17. Olekhnovich, I. & Gussin, G. N. Effects of mutations in the Pseudomonas putida miaA gene: Regulation of the trpE and trpGDC operons in P. putida by attenuation. J. Bacteriol.183, 3256–3260 (2001).

18. J. M. Wang, D., Pierson, L. S., & Pierson, E. A. Disruption of MiaA provides insights into the regulation of phenazine biosynthesis under suboptimal growth conditions in Pseudomonas chlororaphis. Microbiology 163, 94–108 (2017).

19. Tian, Z. X., Fargier, E., Mac Aogáin, M., Adams, C., Wang, Y. P., & O’Gara, F. Transcriptome profiling defines a novel regulon modulated by the LysR-type transcriptional regulator MexT in Pseudomonas aeruginosa. Nucleic Acids Res. 37, 7546–7559 (2009).

20. Ilves, H., Hõrak, R., Teras, R., & Kivisaar, M. IHF is the limiting host factor in transposition of Pseudomonas putida transposon Tn4652 in stationary phase. Mol. Microbiol. 51, 1773–1785 (2004).

21. Mi, J. et al. Investigation of plasmid-induced growth defect in Pseudomonas putida. J. Biotechnol. 231, 167–173 (2016).

22. Lenzen, C., Wynands, B., Otto, M., Bolzenius, J., Mennicken, P., Blank, L. M., & Wierckx, N. High-yield production of 4-hydroxybenzoate from glucose or glycerol by an engineered Pseudomonas taiwanensis VLB120. Front. Bioeng. Biotechnol. 7, 1–17 (2019).

23. Huccetogullari, D., Luo, Z. W., & Lee, S. Y. Metabolic engineering of microorganisms for production of aromatic compounds. Microb. Cell Fact. 18, 1–29 (2019).

24. Furman, B. L. Salicylic acid. Reference Module in Biomedical Sciences, (2018).

25. Vardon, Derek R., et al. cis, cis-Muconic acid: separation and catalysis to bio-adipic acid for nylon-6, 6 polymerization. Green Chem. 18.11, 3397–3413 (2016).

26. Khan, S. A., Chatterjee, S. S., & Kumar, V. Potential anti-stress, anxiolytic and antidepressant like activities of mono-hydroxybenzoic acids and aspirin in rodents: a comparative study. Austin. J. Pharmacol. Ther. 3, 1073 (2015).

27. Kricheldorf, H. R. & Löhden, G. New polymer syntheses, 79. Hyperbranched poly(ester-amide)s based on 3-hydroxybenzoic acid and 3,5-diaminobenzoic acid. Macromol. Chem. Phys. 196, 1839–1854 (1995).

28. Noda, S., Mori, Y., Fujiwara, R., Shirai, T., Tanaka, T., & Kondo, A. Reprogramming Escherichia coli pyruvate-forming reaction towards chorismate derivatives production. Metab. Eng. 67, 1–10 (2021).

29. Wang, J., Zhang, R., Zhang, Y., Yang, Y., Lin, Y., & Yan, Y. Developing a pyruvate-driven metabolic scenario for growth-coupled microbial production. Metab. Eng. 55, 191–200 (2019).

30. Asin-Garcia, E., Batianis, C., Li, Y., Fawcett, J. D., de Jong, I., & Martins dos Santos, V. A. P. Phosphite synthetic auxotrophy as an effective biocontainment strategy for the industrial chassis Pseudomonas putida. Micro. Cell Fact. 21, 156 (2022).

31. Claassens, N. J., He, H., & Bar-Even, A. Synthetic methanol and formate assimilation via modular engineering and selection strategies. Curr. Issues Mol. Biol. 33, 237–248 (2019).

32. Shaw, A. J. et al. Metabolic engineering of microbial competitive advantage for industrial fermentation processes. Science 353, 583–586 (2016).

33. Elmore, J. R., et al. Engineered Pseudomonas putida simultaneously catabolizes five major components of corn stover lignocellulose: Glucose, xylose, arabinose, p-coumaric acid, and acetic acid. Metab. Eng. 62, 62–71 (2020).

34. Damalas, S. G., Batianis, C., Martin-Pascual, M., Lorenzo, V. de, & Martins dos Santos, V. A. P. SEVA 3.1: enabling interoperability of DNA assembly among the SEVA, BioBricks and Type IIS restriction enzyme standards. Microb. Biotech. 13, 1793–1806 (2020).

35. Wirth, N. T., Kozaeva, E., & Nikel, P. I. Accelerated genome engineering of Pseudomonas putida by I-SceI-mediated recombination and CRISPR-Cas9 counterselection. Microb. Biotech. 13, 233–249 (2020).

36. Volke, D.C., Friis, L., Wirth, N.T., Turlin, J., & Nikel, P.I. Synthetic control of plasmid replication enables target- and self-curing of vectors and expedites genome engineering of Pseudomonas putida. Metab. Eng. Commun. 10, e00126 (2020).

37. Asin-Garcia, E., Martin-Pascual, M., Garcia-Morales, L., van Kranenburg, R., & Martins dos Santos, V. A. ReScribe: an unrestrained tool combining multiplex recombineering and minimal-PAM ScCas9 for genome recoding Pseudomonas putida. ACS Synth. Biol. 10, 2672–2688 (2021).

38. Barrick, J. E. et al. Identifying structural variation in haploid microbial genomes from short-read resequencing data using breseq. BMC Genomics 15, 1–17 (2014).

39. Nogales, J., et al. High-quality genome-scale metabolic modelling of Pseudomonas putida highlights its broad metabolic capabilities. Environ. Microb. 22, 255–269 (2020).

40. Nikel, P. I., Kim, J., & de Lorenzo, V. Metabolic and regulatory rearrangements underlying glycerol metabolism in Pseudomonas putida KT2440: Glycerol metabolism in P. putida. Environ. Microb.16, 239–254 (2014).

41. Batianis, C., van Rosmalen, R., Major, M., van Ee, C., Kasiotakis, A., Weusthuis, R. A., & Martins dos Santos, V. A. P. A tunable metabolic valve for precise growth control and increased product formation in Pseudomonas putida. Metab. Eng. 75, 47–57 (2022).

