## Supplementary information for "Shikimate pathway-Dependent Catabolism: enabling near-to-maximum production yield of aromatics"

**Energy generation by wild-type and SDC**

Native and SDC metabolism were simulated with flux balance analysis (FBA) with pyruvate production (EX_pyr_e) as the maximization objective. Then, reduced models were generated containing only predicted active reactions of central carbon metabolism (Supplementary Data 6), as well as exchange reactions for NADH (nadh_c), NADPH (nadph_c), ATP (atp_c), NAD (nad_c), NADP (nadp_c), pyruvate (pyr_c), H^+^ (h_c), H2O (h2o_c), ubiquinone-8 (q8_c), ubiquinol (q8h2_c), oxygen (o2_c), coenzyme A (coa_c), CO2 (co2_c) and inorganic phosphate (pi_c) in the cytoplasm. To find the most efficient overall reaction in terms of ATP generation, pyruvate exchange was set as the objective to maximize, and the bounds of its exchange reaction were constrained to the optimal value. Then, the ATP exchange reaction was set as a new objective to maximize. A glycerol uptake of 1 mmol/gCDW/h was used and the fluxes through the different exchange reactions are defined as coefficients in the overall equations. According to iJN1462 oxidation of NADH via NADH dehydrogenase (reaction NADH16pp) allows the export of 3 H^+^ to the periplasm. The reduction of ubiquinone-8 (q8) to ubiquinol (q8h2), and oxidation of q8h2 by cytochrome oxidase (CYTVo3_4pp) allows the export of 4 H^+^ to the periplasm. Generation of ATP by the ATP synthase (ATPS4rpp) requires the import of 4 H^+^ from the cytoplasm resulting in a yield of 1 ATP per oxidation of 1 q8h2 and 1.75 ATP per oxidation of NADH.

**Overall equation native metabolism (KT2440):**

- *1 Glycerol + 1 Q8 + 1 NAD^+^ + 1 ATP + Pi* 🡪 *1 PYR + 1 Q8H2 + NADH + H^+^ + 2 ATP*

Simplified equation (conversion of reduced cofactors to ATP in the respiratory chain):

- *1 Glycerol* 🡪 *1 PYR + 3.75 ATP*

**Overall equation SDC:**

- *1 Glycerol + 1.8 Q8 + 1.4 NAD^+^ + 1.1 NADPH + 1.1 O_2_ + 2.1 ATP + 1.3 H_2_O 🡪 0.55 PYR + 1.8 Q8H_2_ + 1.4 NADH + 1.4 CO_2_ + 2.1 ADP + 2.1 Pi + 2.9 H^+^*

Simplified equation (assuming conversion of NADH to NADPH and generation of ATP by oxidation of reduced cofactors in the respiratory chain):

- *1 Glycerol + 1.1 O_2_ 🡪 0.55 PYR + 1.4 CO_2_ + 0.23 ATP*

Note: the O_2_ in this equation is not used in the respiratory chain but in the first two reactions converting 4HB into TCA intermediates.

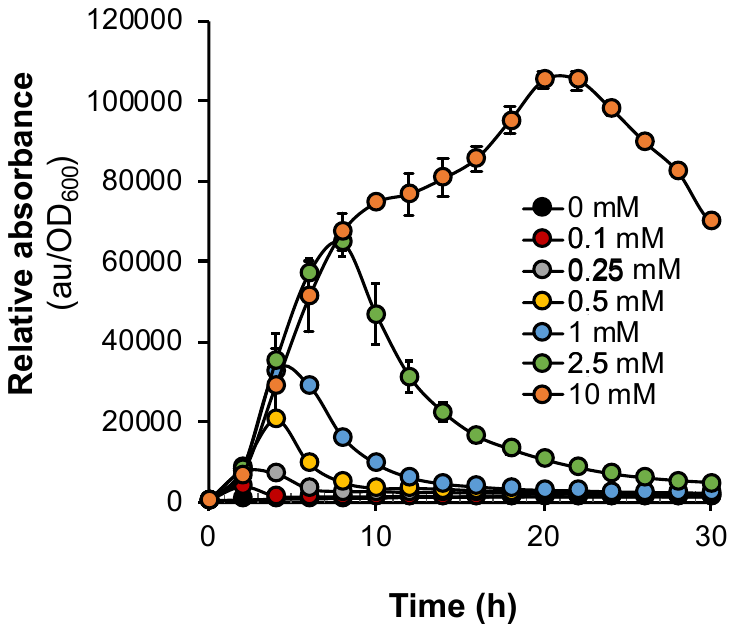

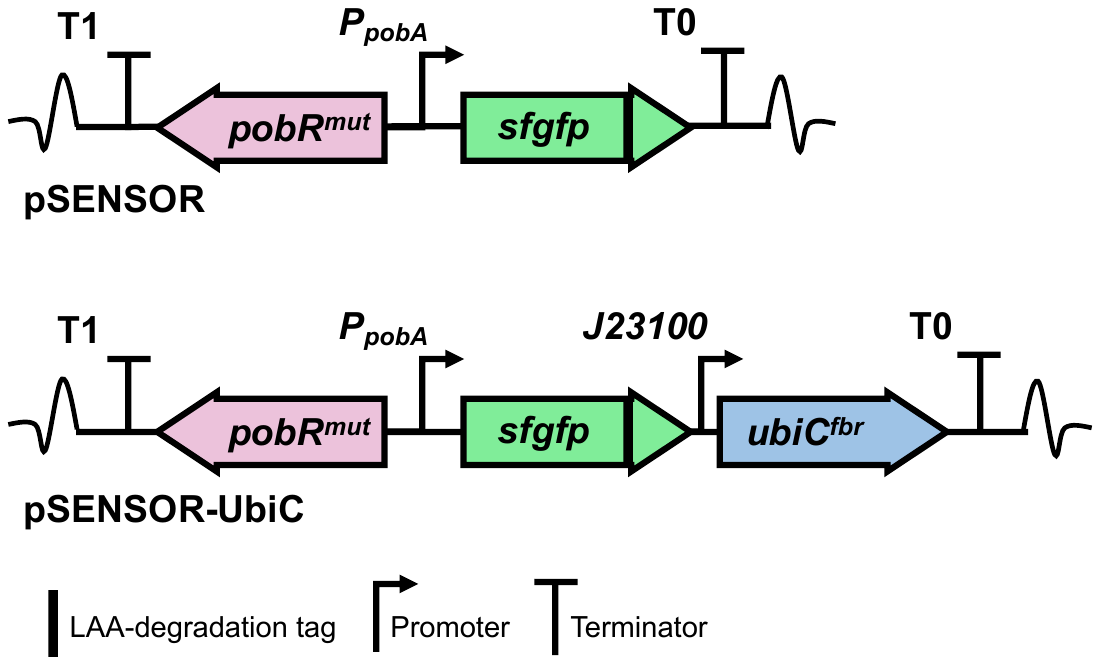

**B**

**Figure 1**. A) Schematic representation of plasmids pSENSOR and pSENSOR_UbiC. B) Relative fluorescence profiles of *P. putida* Δ*glpR* equipped with the 4HB-sensor upon exposure to increasing 4HB levels in glycerol minimal medium. Error bars represent the standard deviation of three biological replicates. Source data are provided as a Source Data file.

**A**

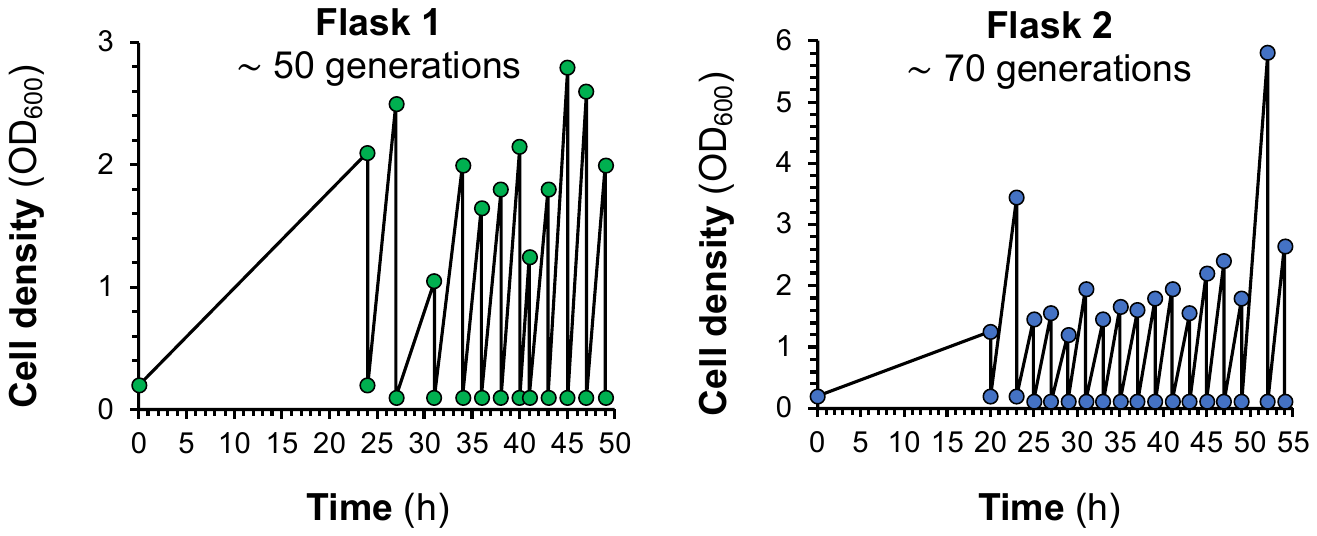

**Figure 2.** Adaptive laboratory evolution (ALE). Growth profiles of the sequential diluted flask cultivations during evolution. Source data are provided as a Source Data file.

**Figure 3**. Screening of evolved mutants after ALE. A) Growth of selected evolved isolates in a microplate reader compared to the parental (ΔPYR) strain and the control (*P. putida* Δ*glpR* pSENSOR_UbiC). B) Relative sfGFP production of the selected evolved isolates on a microplate reader compared to the parental strain and the control. Source data are provided as a Source Data file.

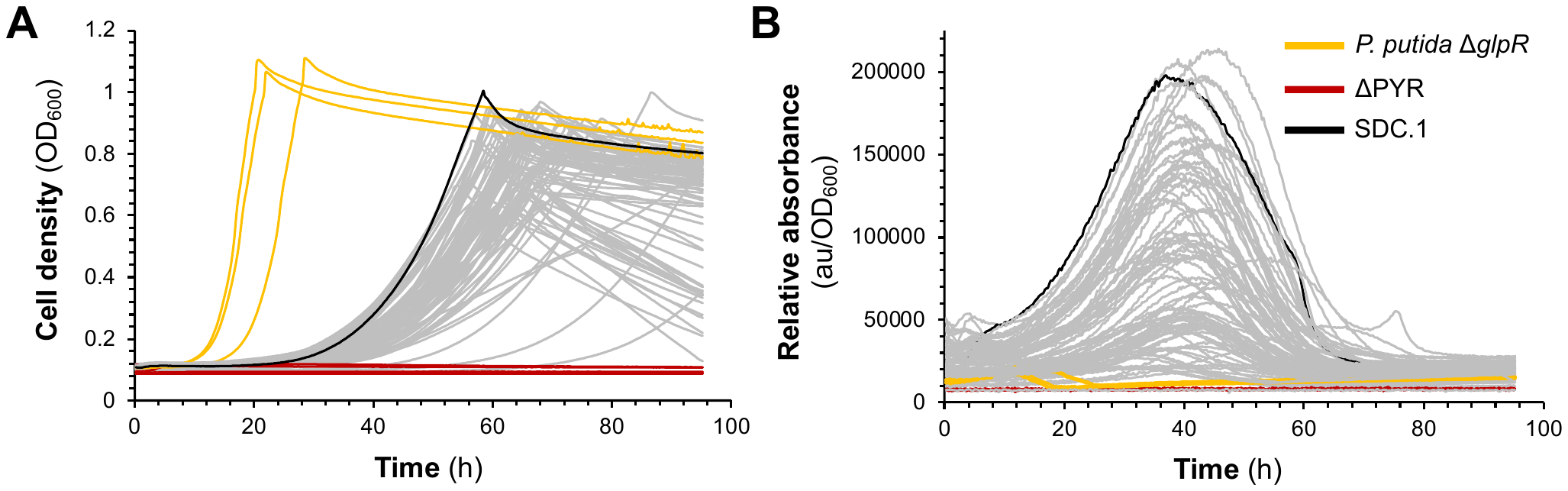

**Figure 4**. A) Identification of the duplicated region from next generation sequencing data, which presented ~2× the number of sequencing reads in SDC.1 and the normal read coverage of the unevolved ΔPYR. The graphs of coverage were generated in Geneious Prime. B) Agarose gel electrophoresis of extracted plasmid pool from strain SDC.1. Lane 1, 1kb Generuler Thermo DNA ladder; Lane 2, plasmid extract.

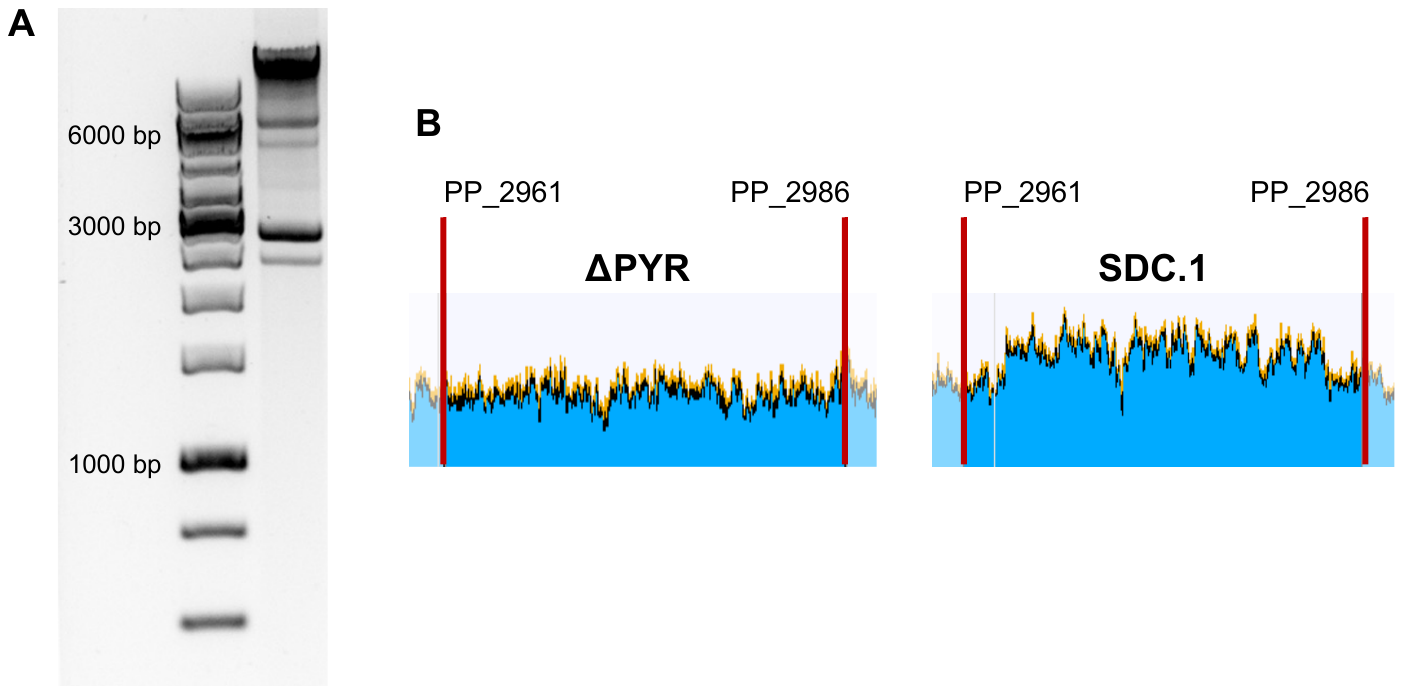

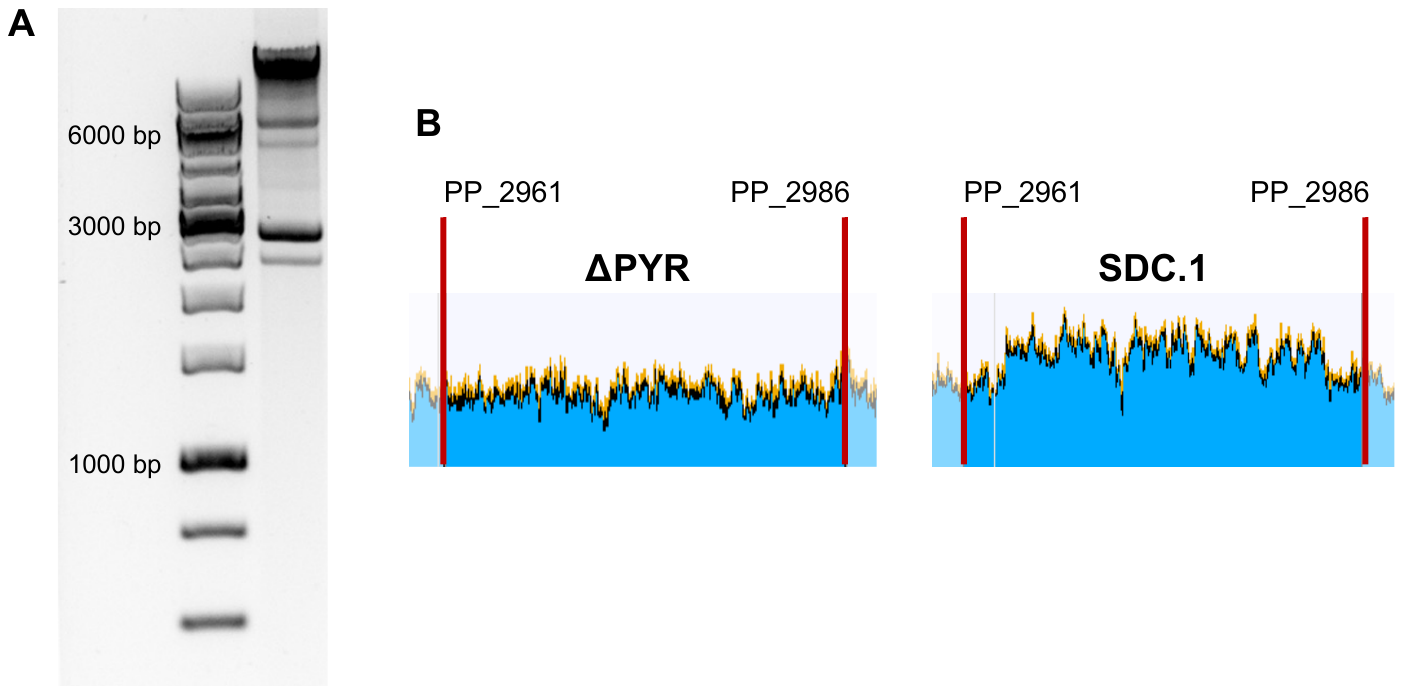

**A**

**B**

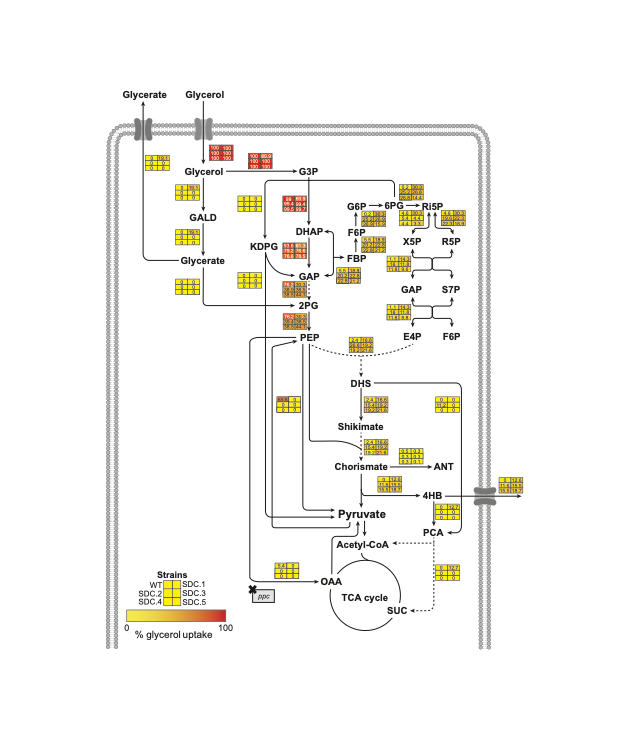

**Figure 5**. Predicted metabolic flux distribution using FBA for the *Pseudomonas putida* KT2440 (WT) and SDC strains. Flux distribution is represented as a percentage of the uptake of 3.95 mmol/gDW/h of glycerol.
